# Sensory cortical ensembles exhibit differential coupling to ripples in distinct hippocampal subregions

**DOI:** 10.1101/2023.03.17.533028

**Authors:** Huijeong Jeong, Vijay Mohan K Namboodiri, Min Whan Jung, Mark L. Andermann

**Author notes:** Correspondence: Vijay Mohan K Namboodiri, Min Whan Jung, Mark L. Andermann.

## Abstract

Cortical neurons activated during recent experiences often reactivate with dorsal hippocampal CA1 sharp-wave ripples (SWRs) during subsequent rest. Less is known about cortical interactions with intermediate hippocampal CA1, whose connectivity, functions, and SWRs differ from those of dorsal CA1. We identified three clusters of visual cortical excitatory neurons that are excited together with either dorsal or intermediate CA1 SWRs, or suppressed before both SWRs. Neurons in each cluster were distributed across primary and higher visual cortices and co-active even in the absence of SWRs. These ensembles exhibited similar visual responses but different coupling to thalamus and pupil-indexed arousal. We observed a consistent activity sequence: (i) suppression of SWR-suppressed cortical neurons, (ii) thalamic silence, and (iii) activation of the cortical ensemble preceding and predicting intermediate CA1 SWRs. We propose that the coordinated dynamics of these ensembles relay visual experiences to distinct hippocampal subregions for incorporation into different cognitive maps.

## Introduction

A key goal in cortical neuroscience is to understand large-scale patterns of activity in recordings from hundreds or thousands of cortical neurons. Numerous studies have indicated that the effective dimensionality of such patterns is most likely significantly lower due to the presence of neuronal ensembles distributed within and across brain areas that exhibit similar sensory tuning properties and strong trial-to-trial correlations in sensory responses (Carrillo-Reid et al., 2015; Clancy et al., 2019; Cossell et al., 2015; Hofer et al., 2011; Ko et al., 2011; Ramesh et al., 2018). Some studies find that neuronal ensembles defined by co-variability of sensory responses can also exhibit co-variability in spontaneous activity during quiet waking (Berkes et al., 2011; Carrillo-Reid et al., 2015; Ch’Ng & Reid, 2010; Han et al., 2008; Hofer et al., 2011; Hoffman & McNaughton, 2002; Kenet et al., 2003; Miller et al., 2014). However, studies in humans argue that ‘default mode’ network patterns of spontaneous cortical activity differ considerably from patterns observed during sensory stimulation (Raichle, 2015). Furthermore, in contrast to cortical ensembles defined by shared co-variability in response to a stimulus, co-variability in spontaneous activity is not usually referenced to specific internally generated events. This has hindered head-to-head comparisons of the nature of neuronal ensembles defined by shared sensory responses and shared “spontaneous” fluctuations.

The hippocampal sharp-wave ripple is an exceptionally well-studied internally generated event (Buzsáki, 2015; ripple for short). Ripples are generated stochastically during immobility or non-REM sleep (Buzsáki, 2015). During waking ripples, the pupil is constricted (McGinley et al., 2015) and the thalamus is strongly suppressed (Logothetis, 2015; Logothetis et al., 2012; Yang et al., 2019), indicating that animals are in a state of particularly low arousal and insensitivity to sensory stimuli. Ripples are often associated with replays, which are temporally compressed sequential reactivations of neurons that were activated during a prior experience. Such replay events are believed to reflect recollection and consolidation of past experiences as well as rehearsal of an internal model (Bhattarai et al., 2020; Carr et al., 2011; Foster, 2017; Gillespie et al., 2021; Gupta et al., 2010; Jung et al., 2018; K Namboodiri & Stuber, 2021; Ólafsdóttir et al., 2018; O’Neill et al., 2010). Consistent with these proposals, ripples are coupled to brain-wide activity changes across cortical and subcortical areas (Logothetis et al., 2012; Karimi Abadchi et al., 2020; Liu et al., 2021; Chrobak & Buzsáki, 1996; Siapas & Wilson, 1998; Jadhav et al., 2016; Rothschild et al., 2017; Gomperts et al., 2015; Pennartz et al., 2004). Notably, ripples do not always occur over the entire longitudinal axis of the hippocampus; instead, localized ripples are frequently observed (De Filippo & Schmitz, 2022; Nitzan et al., 2022; Patel et al., 2013; Sosa et al., 2020). These localized ripples have different features such as amplitude, duration, and frequency (Nitzan et al., 2022), and recruit distinct populations of nucleus accumbens neurons (Sosa et al., 2020). In addition, different parts of the hippocampus serve distinct functions with distinct anatomical connectivity with the neocortex (Fanselow & Dong, 2010; Moser & Moser, 1998; Strange et al., 2014). Thus, we reasoned that localized hippocampal ripples may couple with distinct networks of cortical neurons, and that this coupling may define previously unknown functional cortical ensembles.

To test this hypothesis, we analyzed the activity of hundreds of neurons recorded simultaneously from various regions of the visual cortex, hippocampus, and thalamus (Neuropixels data, Functional Connectivity dataset, Allen Institute; Siegle et al., 2021). We identified three distinct ripple-associated cortical activity patterns: two cortical clusters activated in conjunction with dorsal or intermediate CA1 ripples, and one suppressed before either type of ripple event. Neurons in the same cluster exhibited correlated spiking across arousal levels, especially during low arousal, even in the absence of ripple events. Neurons in each cluster were distributed nearly evenly across primary and higher visual cortical areas and exhibited similar visual response properties. These findings suggest that duplicate copies of visual information are routed to different regions of hippocampus via distinct sensory cortical ensembles. This process may facilitate the integration of visual experiences into multiple brain-wide networks and the updating of distinct internal models during quiet waking.

## Results

### Three distinct patterns of ripple-associated cortical neuronal activity

Spiking activity was measured simultaneously from six Neuropixels probes during passive presentation of various visual stimuli in awake mice (Allen Institute, the Functional Connectivity dataset; Fig. 1A). Each session also included a thirty-minute ‘spontaneous period’, during which a static gray screen was shown to the mice. Each Neuropixels probe targeted one of six visual cortical areas and passed through the dorsal (dCA1) or intermediate CA1 (iCA1) (Fig. 1B). We focused on the analysis of hippocampal ripples and associated cortical activity during the spontaneous period. As reported previously (De Filippo & Schmitz, 2022; Nitzan et al., 2022; Patel et al., 2013; Sosa et al., 2020), we observed some ripples propagating along the longitudinal axis of CA1 and others localized to dCA1 or iCA1 (Fig. 1C-E). Ripples were classified into global (ripples detected at both dCA1 and iCA1 with temporal overlap; 47.3±14.6% and 58.6±17.3% of all ripples detected at dCA1 and iCA1, respectively), dCA1 (ripples detected only at dCA1), and iCA1 (ripples detected only at iCA1). Ripple strength did not differ significantly between the dCA1 and iCA1, but ripple duration was longer in dCA1 (41.2 ± 5 ms) than iCA1 (32.1 ± 5 ms, p<0.001). Also, global ripples were stronger and lasted longer than local ripples, with no consistent bias in travel direction (Fig. S1).

**Fig. 1.**
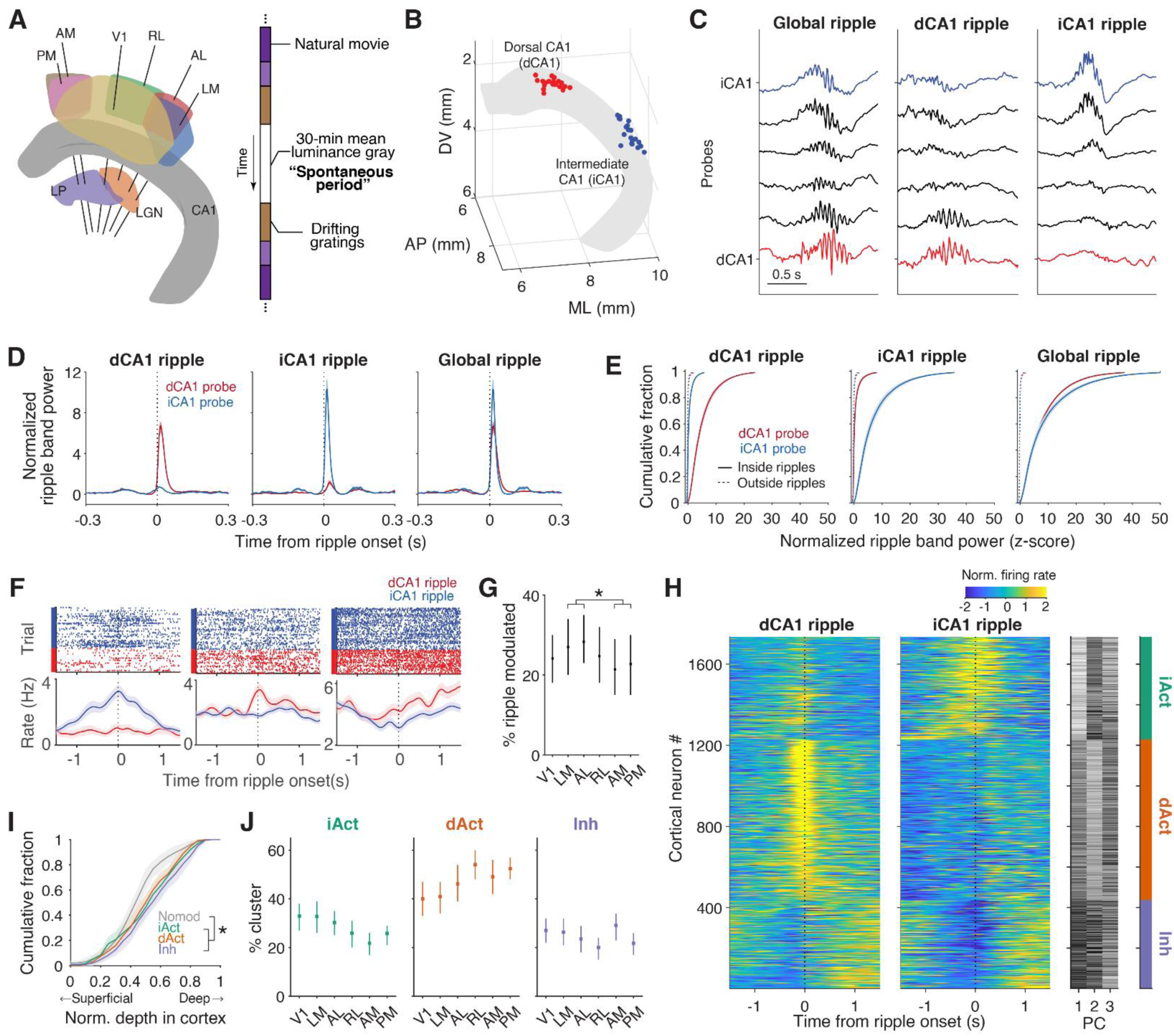
Hippocampus ripples classify visual cortical neurons into four types. **A.** Neuropixels probe locations and recording session timeline. Neuropixels probes targeted six visual cortical areas and also passed through the hippocampus and thalamus. Animals passively viewed various visual stimuli including natural movies and drifting gratings. Most analyses used data from a thirty-minute spontaneous period during which mean luminance gray was presented. **B.** CA1 pyramidal layer recording locations (n=20 sessions). Red and blue dots represent the most medially and laterally positioned probes in each session, which target dorsal and intermediate CA1, respectively. **C.** Three representative CA1 local field potential (LFP) traces from 6 Neuropixels probes showing ripple events. Red and blue traces represent LFP from dorsal (dCA1) and intermediate (iCA1) CA1, respectively. Some ripples span dorsal and intermediate CA1 (global ripples; left), while others are more localized in dCA1 (middle) or iCA1 (right). **D.** Normalized ripple band (100-250 Hz) power from dCA1- and iCA1-targeting probes around different types of ripples. **E.** Cumulative fraction of normalized ripple band power at time bins with (Inside ripples) or without ripples (Outside ripples). Each graph was plotted until 0.99 cumulative fraction (See Methods). **F.** Three sample visual cortical neuron responses to dorsal (red) and intermediate (blue) ripples. Top, spike raster plots; bottom, spike density functions. **G.** Fractions of dCA1 or iCA1 ripple-responsive regular spiking neurons. Ripples modulated lateral visual cortical neurons (LM and AL, 32.2±16.5%) more than medial ones (AM and PM, 27.4 ± 15.4 %; n=17 sessions targeting both areas). **H.** Ripple-responsive visual cortex neurons were clustered into three groups based on average activities around ripples (see Methods). The first three principal components of average ripple responses are shown in gray scale. Green, orange and purple colors indicate iAct, dAct, and Inh clusters, respectively. **I.** Cumulative fraction of normalized neuron depth in each cluster (0, surface; 1, white matter). Ripple non-responsive neurons were located more superficially than ripple-responsive neurons, but the depth of ripple-responsive neurons did not vary across clusters. **J.** Fractions of cluster-specific ripple-modulated neurons per area. Note neurons in every cluster are widely distributed across visual areas, with slight biases of iAct and dAct neurons towards lateral and medial visual areas, respectively. Error bars in all figures indicate mean±95% bootstrap confidence intervals unless noted. See Table S1 for detailed results of statistical tests.

We next examined the activity of visual cortical neurons during ripples. We found that 30.4 ± 14.6% of regular spiking neurons changed their activity significantly around ripples (Fig. 1F; See Methods). All visual cortical areas, including the primary visual cortex, had similar proportions of ripple-modulated neurons (∼20-30%), but the fraction was slightly larger in the lateral (LM and AL) than medial visual cortical areas (AM and PM; Fig. 1G). Notably, many neurons were modulated preferentially by either dCA1 or iCA1 ripples (Fig. 1F, H). We classified ripple-modulated cortical neurons based on their activity profiles around dCA1 and iCA1 ripples using principal component analysis (PCA) and k-means clustering (Fig. S2). We identified three clusters of ripple-modulated neurons: two showed preferential activation around either iCA1 (iAct) or dCA1 ripples (dAct), and one showed suppression around both types of ripples (Inh; Fig. 1H). The dAct and iAct clusters increased activity, while the Inh cluster decreased activity around the global ripple (Fig. S1C). The activity profiles of these three clusters matched those of the first three PCs, and together they explained ∼56% of the total variance in neural activity during the 3-s time window centered around ripple onset (Fig. S2B).

When compared to ripple-non-modulated neurons (Nomod, 69.6 ± 14.6%), ripple-modulated neurons were more likely to be in deeper layers (Fig. 1I). In particular, ripple-modulated neurons were less likely to be in putative layer 4 and more likely to be in putative layer 6 than non-modulated neurons (Fig. S3). This suggests that ripple-modulated neurons are more likely to send feedback to the thalamus than to receive thalamic feedforward sensory inputs (Jia et al., 2022; Lamme et al., 1998). Although neurons from each visual cortical area were included in all three ripple-modulated clusters, the iAct and dAct clusters were biased towards the lateral and medial visual cortices, respectively (Fig. 1J). These regional biases were more pronounced in the analysis of local field potentials (Fig. S4). This may reflect stronger connections between the medial visual cortices and dorsal hippocampus via the retrosplenial cortex, and between the lateral visual cortices and intermediate hippocampus via the entorhinal and temporal association cortices (Cenquizca & Swanson, 2007; Meier et al., 2021; Wang et al., 2012; Yao et al., 2021).

### Ripple-modulated cortical clusters are functional ensembles spanning the visual hierarchy

It has been suggested that an ensemble of co-active neurons may be a functional unit of neural computation (Yuste, 2015). We measured neuronal activity correlations in the absence of a ripple (ripple epochs were removed; see Methods) during the spontaneous period to see if ripple-modulated cortical clusters represent functional ensembles (Fig. 2A). On average, pairs of ripple-modulated neurons had significantly stronger correlations than non-modulated neurons (Fig. 2B). Also, pairs of neurons from the same cluster showed stronger correlations than those from different clusters (Fig. 2B). Furthermore, regardless of whether two neurons were in the same or different visual areas, activity correlations were similarly high for those in the same cluster, and similarly low for those in different clusters (Fig. 2C). This was similarly observed when analysis was separately performed for each cluster, but with strongest area dependency in dAct cluster (Fig. S5A). dAct neurons showed high within-cluster correlation in each visual cortical area (Fig. S5B), but lower between-area correlation. Collectively, these results suggest that ripple-modulated cortical clusters, especially iAct and Inh clusters, may represent coherent functional ensembles spanning multiple visual cortical areas, while dAct cluster is composed of a collection of ensembles within individual visual areas.

**Fig. 2.**
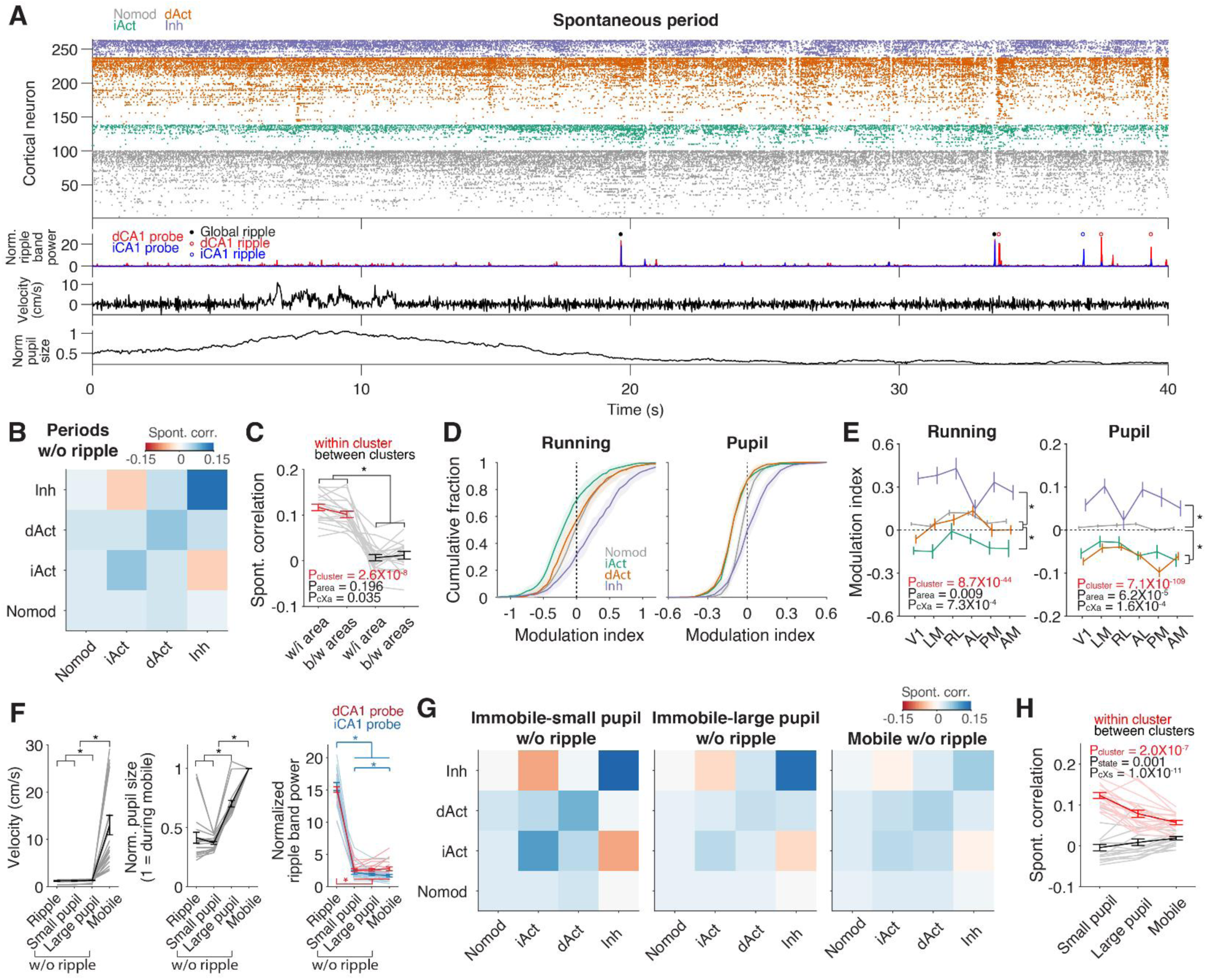
Visual cortex clusters maintain correlated activity outside ripple events. **A.** Spike raster plots of all neurons during an example spontaneous period. Normalized ripple band power, running speed, and pupil size are also shown. Pupil size was normalized by average pupil size during running (> 2 cm/s). **B.** Mean neuronal correlations within and across clusters during the spontaneous period. We excluded ripple-containing 2-s time bins. The heatmap was averaged across sessions (n = 18; only sessions with at least one cell in each cluster were used). The correlation within a ripple-modulated cluster (diagonal line; 0.09±0.006, averaged across the three ripple-modulated clusters) is stronger than that between clusters (0.01±0.007, averaged across iAct-dAct, iAct-Inh, and dAct-Inh) or that within the Nomod group (0.02±0.003). **C.** Mean spontaneous neuronal correlations within (w/i area) and between areas (b/w areas). Neuronal pairs within a cluster have a higher spontaneous correlation than those between clusters in both cases. **D.** Cumulative fraction of running (left) or pupil (right) modulation index. Positive modulation index indicates neuronal activity is higher during high-arousal (running or large pupil) than low-arousal (stationary or small pupil). To avoid potential confounding effects of running, pupil modulation index was calculated using only immobile-state data. **E.** Average cluster-specific running and pupil modulation indices. **F.** Average running speed, normalized pupil size, and normalized ripple band power in four states: a ripple-containing state (Ripple), two immobile states with small pupil (Small pupil) or large pupil (Large pupil), and a mobile state (Mobile). Ripple-containing periods were excluded for the latter three states. Normalized ripple band power used peak power rather than average power within each 2-s analysis window. Despite clear differences in running speed and pupil size, ripple power was minimally different across the latter three states. **G.** Average neuronal correlations in three behavioral states. **I.** Average correlation of within-or between-clusters neuronal pairs during three behavioral states. Within-cluster correlation is significantly greater than between-cluster correlation even in the mobile state, though the difference is smaller than in immobile states.

Because ripples occur at low arousal, we examined how arousal level influences the activity of ripple-modulated neurons using running speed and pupil size as proxy for arousal level (Reimer et al., 2014; Vinck et al., 2015) (Fig. 2A). We found that the activity of the Inh and iAct clusters are positively and negatively correlated, respectively, with running speed and pupil size (Fig. 2D). This was true regardless of neuronal location (Fig. 2E). As strong within-cluster correlations can be explained by shared state-dependent modulation of neural activity, we divided the spontaneous period into three states based on running speed and pupil size (excluding ripple epochs): immobile with small pupil dilation (Small pupil), immobile with large pupil dilation (Large pupil), and mobile (Mobile; Fig. 2F). Because ripple epochs were excluded, ripple power did not vary significantly across the three states (Fig. 2F). Even though the analysis was limited to a narrow range of arousal level and ripple power was similarly low in all three states, within-cluster neuronal correlations remained higher than between-cluster correlations at each arousal level (Fig. 2G, H). We observed similar pattern when separately analyzing each ensemble (Fig. S5C). These results indicate that strong within-cluster correlations cannot be fully accounted for by shared state-dependent modulation. Strong within-cluster correlations outside ripple epochs are more likely to be caused by other mechanisms such as strong recurrent connectivity within each cluster (Ko et al., 2011).

We also tested the possibility that our results were influenced by subthreshold ripple-like events. In contrast to this possibility, we obtained similar results even when we limited our analysis to time periods with the lowest 20% of ripple band power in the small pupil state (Fig. S6A, B). We additionally found that as arousal level decreases, the difference between within-cluster and between-cluster correlations becomes stronger (Fig. 2H). This suggests that these spontaneous ensembles might be specialized to function during low-arousal states, when external sensory influences are minimal.

### Visual tuning properties are similar across spontaneous ensembles

Recent studies using large-scale recording techniques showed that spontaneous and sensory-evoked activity are nearly orthogonal across species (Avitan et al., 2021; Liu et al., 2021; McGuire et al., 2022; Stringer et al., 2019; Triplett et al., 2020; but see Berkes et al., 2011; Han et al., 2008; Hoffman & McNaughton, 2002). We reasoned that the Inh cluster, which is suppressed during ripples but active during high arousal (Fig. 2D), may be more sensitive to sensory stimuli than the clusters that are activated during ripples (dAct and iAct clusters). To determine whether visual responses vary across ripple-modulated clusters, we analyzed visual responses of neurons with a significant receptive field (Fig. 3A). The proportion of neurons with a receptive field was higher in Nomod neurons than ripple-modulated neurons (consistent with the finding that Nomod neurons are prevalent in layer 4), but did not vary significantly across ripple-modulated clusters, contrary to our prediction. Furthermore, the size of the receptive field did not differ significantly across clusters (Fig. 3A).

**Fig. 3.**
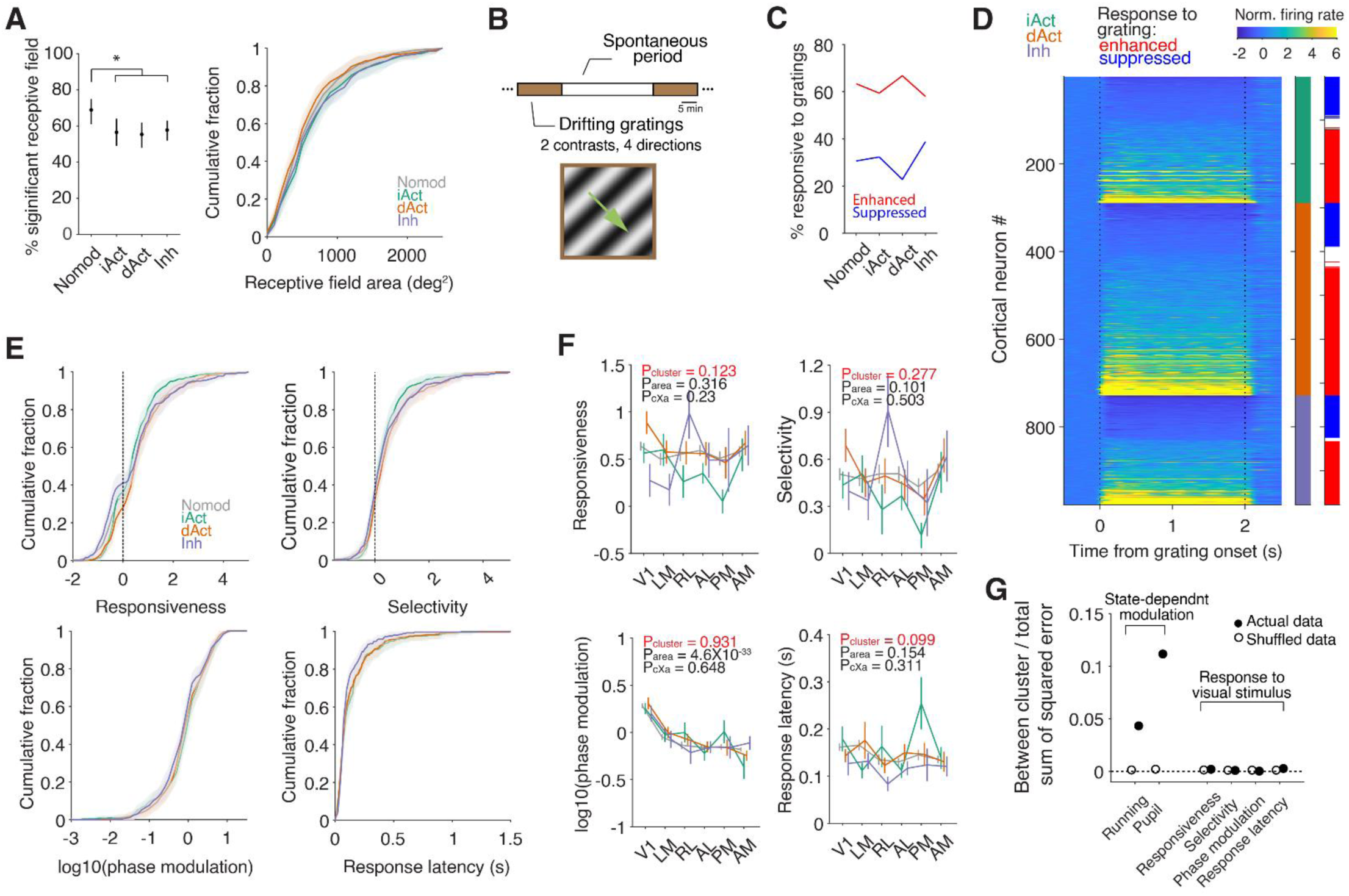
Visual properties did not differ across ripple-coupled ensembles. **A.** Fraction of neurons with significant receptive fields in each cluster (left) and cumulative distribution of receptive field size (right). Only neurons with significant receptive fields were used for the remaining analyses in Fig. 3. Nomod neurons were more likely to have significant receptive fields than ripple-modulated clusters. Receptive field size did not vary significantly across clusters. **B.** Schematic of session structure. Visual response characteristics were determined by analyzing neural responses to drifting gratings in Fig. 3. **C.** Fractions of grating-responsive neurons. **D.** Responses of ripple-modulate neurons to drifting gratings. Neurons were sorted by cluster identity and grating response. Each ensemble had neurons with enhanced and suppressed responses to drifting gratings. **E.** Cumulative fraction of neurons as a function of responsiveness to the preferred direction, selectivity between the preferred and orthogonal directions, phase modulation, and response latency. **F.** Cluster responsiveness, selectivity, phase modulation, and response latency averaged by area. Note overall similarity in visual response properties across clusters. **G.** Explained neural activity variance by behavior states and visual response properties. Open circle denotes explained variance from shuffled cluster identities.

We then compared neuronal responses to drifting gratings which were presented during blocks of trials that immediately preceded or followed the spontaneous recording period (Fig. 3B). Some neurons in each cluster increased their activity, while others decreased their activity in response to the grating stimulus (Fig. 3C, D). We measured responsiveness (activity in the presence versus absence of a stimulus), selectivity (response to preferred versus orthogonal directions), phase modulation, and response latency. Neurons in each cluster had a wide range of values for each tuning property, but these distributions were largely similar across clusters (Fig. 3E). The variability in neuronal tuning properties could not be explained by cluster identity, even after accounting for regional differences (Fig. 3F). While each neuron’s cluster identity could explain more than 10% of the variance in pupil size-related modulation and ∼4% of the variance in running speed-related modulation (cf. Fig. 2D), cluster identity barely explained any variance in visual tuning (Fig. 3G). In sum, there was no difference in the distribution of visual response properties across ripple-modulated and non-modulated clusters.

### Coordinated activity of cortical ensembles and thalamic neurons

The thalamus acts as a state-dependent gate for sensory inputs to the cortex (Aydın et al., 2018; Busse, 2018; Erisken et al., 2014; Liang et al., 2020; Molnár et al., 2021; Reinhold et al., 2023; Spacek et al., 2022; Williamson et al., 2015). Thalamic activity decreases around hippocampal ripples (Logothetis et al., 2012; Logothetis, 2015; Lara-Vásquez et al., 2016; Yang et al., 2019; Chambers et al., 2022), potentially preventing sensory stimuli from interfering with ripple-related neural processing. We took advantage of the simultaneous recordings from the hippocampus, cortex, and thalamus to examine the relationships between ripple-modulated clusters and thalamic activity. Neurons in different thalamic nuclei showed highly synchronous activity during the spontaneous period (Fig. S6C, D). Inh and iAct clusters showed strong positive and negative correlations with thalamic activity, respectively, while neither dAct cluster nor Nomod neurons showed consistent coupling to thalamic activity (Fig. 4A, B). This selective coupling between the thalamus and specific cortical clusters was strongest in the lowest arousal state (Fig. 4B, C). This pattern of correlations did not reflect direct connectivity between specific thalamic and cortical areas (Fig. S7). Thus, the observed coupling between the thalamus and visual cortical clusters likely depends more on a shared modulation mechanism (e.g., via neuromodulators [S.-H. Lee & Dan, 2012] and/or a hub region such as the thalamic reticular nucleus [Takata, 2020]) than it does on direct anatomical connectivity.

**Fig. 4.**
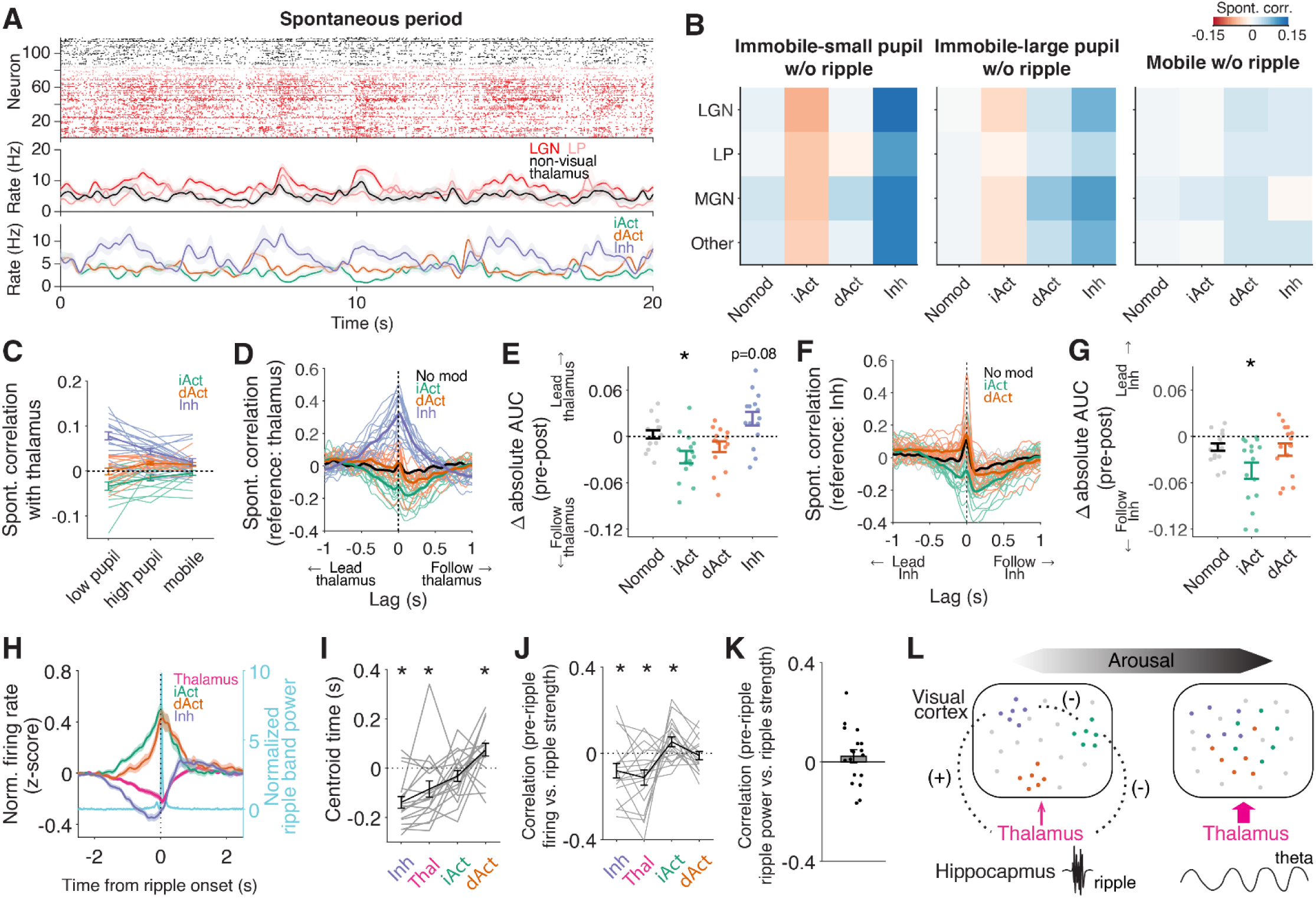
Coordinated visual cortical and thalamic activity precedes a ripple during the low arousal state. **A.** Top: Raster plots of neuronal activity in visual (lateral geniculate nucleus, LGN; lateral posterior nucleus, LP) and non-visual thalamic areas (medial geniculate nucleus, MGN; other thalamic areas, other). Bottom: Mean ±s.e.m. (shading) of simultaneously recorded neuronal activity in each thalamic or cortical group (for single-neuron data from cortex, see last 20 s of Fig. 2A). **B.** Average noise correlation across all pairs within or between clusters in visual cortex and thalamus (cf. Fig. 2G). **C.** Average spontaneous cross-correlation between each cluster and thalamus (including both visual and non-visual thalamic nuclei). Correlation between each cluster and thalamus increases as arousal level decreases. Each line indicates an individual session. **D.** Cross-correlation between thalamus and each cortical cluster during immobility with small pupil size. iAct and Inh ensembles showed positive and negative correlations with thalamus, respectively, while dAct neurons and NoMod showed weaker correlations. Thalamic activity precedes (or follows) cortical cluster activity when biases are positive (or negative). **E.** The difference in absolute area under the cross-correlation curve (Δabsolute AUC) between before (pre; 0.5 s) and after (post; 0.5 s) time zero. Activation of the Inh ensemble leads the activation of thalamus, which is followed by suppression of iAct ensemble. Thus, suppression of Inh neurons inhibits thalamic neurons and disinhibits iAct neurons. **F-G.** Same format as **D-E**, but between the Inh ensemble and other ensembles. Activation of the Inh ensemble leads suppression of the iAct ensemble. *, paired t test with Nomod. **H.** The average firing rate of visual cortex ensembles and the thalamus around global ripple onset. Normalized hippocampal ripple band power is also shown. **I.** Centroid time of ripple-associated activity of each cluster (in the ±1s surrounding ripple onset). Thalamic and visual cortical activity around hippocampal ripples change in a predictable temporal sequence. **J.** Correlation between neural activity 1-s prior to ripple onset and ripple strength. **K.** Correlation between hippocampal ripple band power 1-s prior to ripple onset and ripple strength. **L.** Schematic illustrating how arousal affects visual cortex ensemble-thalamus interactions. Each dot is a visual cortical neuron with its color representing cluster identity (Green, iAct; Orange, dAct; Purple, Inh). Dashed lines connect groups with significant activity correlations, with ‘+’ or ‘-‘ indicating the sign of correlation. Pink arrows indicate overall thalamic activity across arousal levels. The strength of within-ensemble correlations is represented by the proximity of dots. The predominant oscillatory event in the hippocampus at each arousal level (ripple or theta oscillation) is also shown.

Next, we performed a cross-correlation analysis to investigate temporal ordering of thalamic and cortical neuronal activity in the absence of hippocampal ripples. This analysis was restricted to the small pupil, immobile state during which thalamocortical correlations were strongest. Inh cluster activity preceded and was positively correlated with thalamic activity. In contrast, iAct cluster activity followed and was negatively correlated with thalamic activity (Fig. 4D, E). These findings raise the possibility that, even in the absence of hippocampal ripples, decreased activity of Inh cluster neurons—some of which are located in deep cortical layers (Fig. 1I) and likely project to thalamus—may deactivate thalamic neurons, which in turn facilitates iAct cluster activity.

To further test this idea, we examined the temporal relationship between cortical ensembles, focusing on Inh-iAct cluster pairs which show anti-correlated spontaneous activity (Fig. 2B). This analysis yielded consistent results. The suppression of Inh cluster preceded the activation of iAct cluster (Fig. 4F, G). This temporal relationship was also observed prior to the onset of a ripple—Inh cluster activity was suppressed before the suppression of thalamic activity, which was followed by the activation of iAct cluster (Fig. 4H, I; Fig. S1D, E for dCA1 and iCA1 ripples). Importantly, the strength of the upcoming ripple was significantly correlated with the levels of thalamic activity suppression, Inh cluster activity suppression, and iAct cluster activity enhancement during the 1-s period prior to ripple onset (Fig. 4J). These correlations were not merely due to the influence of sub-threshold ripples, as ripple-band power during the same 1-s period was not related to subsequent ripple strength (Fig. 4K). Collectively, these results suggest that spontaneous cortical ensembles may play a role in biasing the brain toward internal processing by suppressing external sensory processing of thalamocortical neurons (Fig. 4L).

## Discussion

Understanding hippocampal-cortical interactions is important to determining how the brain builds and uses cognitive maps (K Namboodiri & Stuber, 2021; Whittington et al., 2022). Despite well-known functional variations along the hippocampal longitudinal axis (Fanselow & Dong, 2010; Moser & Moser, 1998; Strange et al., 2014), little is known about how different hippocampal regions interact with the cortex. According to Sosa et al. (2020), ripples that occur selectively in the dCA1 or ventral CA1 (vCA1; iCA1 is intermediate between dCA1 and iCA1) recruit different populations of nucleus accumbens neurons, and neurons coupled to ripples in each of these regions are modulated differently by novelty and reward. This suggests that distinct brain-wide networks with different functions are linked to different hippocampal subregions. In the present study, we identified two distinct visual cortical ensembles that are activated around ripples in dCA1 or iCA1 (dAct and iAct ensembles, respectively). The dCA1 and iCA1 ripples differed in duration, magnitude, peak frequency, and pre-ripple thalamic suppression (Nitzan et al., 2022; Fig. S1); their coupled cortical ensembles also differed in modulation by arousal state and coupling with thalamus and with the Inh ensemble. Considering the differences in anatomical connectivity, molecular composition, and functional roles of dCA1 and iCA1 (Fanselow & Dong, 2010; Jarzebowski et al., 2022; Jin & Lee, 2021; Moser & Moser, 1998; Strange et al., 2014), the iAct and dAct ensembles may be part of separate brain-wide networks that serve related but distinct functions.

Proposed functional variations along the hippocampal longitudinal axis include cognitive versus affective signal processing (Fanselow & Dong, 2010; Moser & Moser, 1998; Strange et al., 2014), fine-grained versus coarse representation of information (Harland et al., 2017; Jung et al., 1994; Poppenk et al., 2013), and faithful versus subjective representation of experienced events (Biane et al., 2022; Yun et al., 2023). Thus, it is likely that dAct and iAct ensembles have different functions. Consistent with this, the centroid of dAct activity followed ripples (global ripple, 0.07±0.03 s; dCA1 ripple, 0.02±0.02) while that of iAct activity preceded ripples on average (global ripple, -0.03±0.02 s; iCA1 ripple, -0.06±0.02 s; Fig. 4I, Fig. S1E). Since it has been suggested that post-ripple cortical activation might reflect memory consolidation as it often accompanies a volley of ripples (Karimi Abadchi et al., 2020), dAct activity may contribute more to the consolidation of previous experiences. On the other hand, iAct ensemble predicts the strength of upcoming ripples (Fig. 4J) and might therefore have some causal influence on ripples. Further, activities of iCA1 ripple-coupled cortical ensembles (iAct and Inh) are more coherently modulated by arousal than dAct ensemble or Nomod neurons (Fig. 2D,E). This may be an indication that iCA1-linked neurons are more sensitive to internal states than dCA1-linked neurons. In addition, the dAct ensemble showed higher spontaneous correlations for neuron pairs within than between areas (Fig. S5A) with high within-cluster correlation in each area (Fig. S5B). This suggests the existence of multiple dCA1 ripple-coupled sub-ensembles, each locally enriched within a given visual area. Meanwhile, pairwise spontaneous correlations within the iAct or Inh ensembles were similar regardless of whether the two neurons were within the same or different areas (Fig. S5A). Collectively, we speculate that multiple brain-wide networks coupled to different hippocampal areas may construct distinct cognitive maps: the dCA1-coupled network may represent fine-grained cognitive maps faithfully reflecting the external environment by virtue of connectivity to distinct fine-grained cortical areas. In contrast, the iCA1-coupled network may represent coarse-grained cognitive maps emphasizing arousal-inducing features of the external environment by virtue of connectivity to a broad swathe of cortical areas.

All three spontaneous cortical ensembles spanned all visual cortical areas examined in this study. Given that different visual cortical areas process different types of visual information (Andermann et al., 2011; Marshel et al., 2011), the function and co-activity of visual cortical neurons are determined in part by their anatomical locations. However, spontaneous correlations were similarly high within a ripple-coupled ensemble for within-area neuron pairs and across-area pairs, particularly for iAct and Inh ensembles. This lack of correlation between ripple-related ensemble identity and spatial location suggests that ripple-related ensembles are organized orthogonal to visual function. Consistent with this, the distribution of visual response profiles was very similar across ensembles despite significant differences in the degree and nature of ripple-related activity. This suggests that multiple copies of visual information are routed during rest to distinct large-scale neural networks linked to different hippocampal areas. It is also worth noting that neurons in the primary visual cortex showed comparable rates of ripple coupling as neurons in higher visual areas (Fig. 1G, J), which may explain the presence of spatial coding in the primary visual cortex (Fiser et al., 2016; Flossmann & Rochefort, 2021; Haggerty & Ji, 2015; Ji & Wilson, 2007; Saleem et al., 2018).

Before a ripple, we observed the following sequence: Inh ensemble suppression, thalamic silence, and iAct ensemble activation. What might be the identity of Inh ensemble neurons in visual cortex? Deep layer cortical neurons send strong inputs to the thalamus (Thomson, 2010), and their suppression reduces state-dependent modulation of thalamic activity (Molnár et al., 2021; Reinhold et al., 2023; but see Nestvogel & McCormick, 2022). Similar proportions of layer-6 corticothalamic neurons in the primary visual cortex increase or decrease their activity in response to visual stimuli, and are positively modulated by arousal regardless of response direction (Augustinaite & Kuhn, 2020). Thus, the overall activity pattern of lnh neurons is similar to those of layer-6 corticothalamic neurons. Thalamic suppression may therefore result from the suppression of the corticothalamic subset of Inh ensemble neurons prior to ripple onset (Reinhold et al., 2023). Additionally, local inhibitory cortical interneurons may mediate the disinhibition of the iAct ensemble following the suppression of the Inh ensemble (Bortone et al., 2014). These events may precede ripples to prepare hippocampus-coupled brain-wide networks for internal, rather than external, signal processing. Collectively, these results significantly enhance our understanding of hippocampo-cortico-thalamic interactions during internal and external signal processing.

## Acknowledgements

We thank L. Frank, N. Nguyen, and the Namboodiri, Jung and Andermann labs for useful feedback. This project was supported by NIH R00MH118422, R01MH129582, R01AA029661, and a Scott Alan Myers Endowed Professorship (V.M.K.N.), the Research Center Program of the Institute for Basic Science (IBS-R002-A1; M.W.J.), and NIH DP2 DK105570, DP1 AT010971-02S1, R01 MH12343, a McKnight Scholar Award, a Harvard Mind Brain Behavior Interfaculty Initiative Faculty Research Award, and the Harvard Brain Science Initiative Bipolar Disorder Seed Grant with support from Kent and Liz Dauten (M.L.A.).

## Author contributions

H.J., M.W.J. and M.L.A. conceived the project. H.J. conducted all analyses with feedback from V.M.K.N., M.W.J. and M.L.A. H.J., V.M.K.N., M.W.J. and M.L.A. wrote the manuscript.

## Supplementary Materials for

This PDF file includes:

Figs. S1 to S7

Methods

Table. S1 (statistical results)

**Fig. S1.**
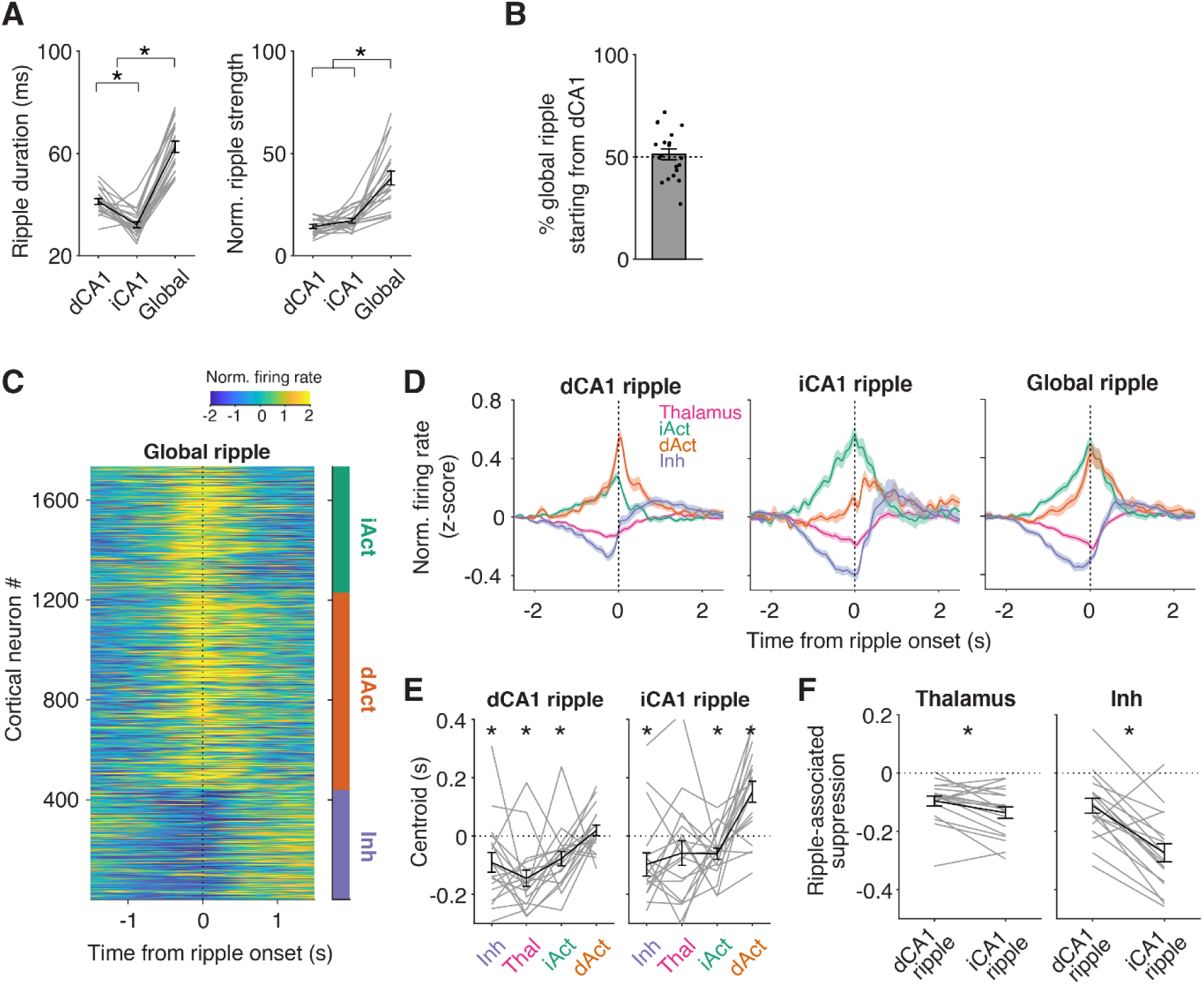
Properties of different ripple types. **A.** Ripple duration (left) and strength (right) of three different ripple types. The duration of global ripples was measured as time from onset of the first detected ripple (either from dCA1 or iCA1) until ripple band power in both dCA1 and iCA1 goes below a threshold of 2 standard deviations above mean ripple power. The strength of global ripples was measured as the maximum power of simultaneously detected ripples from dCA1 and iCA1. Gray lines: individual sessions (n=20, one session per mouse). Black line and errorbars: mean ± s.e.m. across sessions. **B.** Fraction of global ripples that initially arise from dCA1 and travel toward iCA1. dCA1 ripples did not consistently lead or lag iCA1 ripples. **C.** Activity profile of visual cortical neurons around global ripples. Neurons are ordered by cluster identity as in Fig. 1H. **D.** Average activity of visual cortical clusters and thalamus around different ripple types. The traces surrounding global ripples are the same as in Fig. 4H. Inhibition of activity in the thalamus and in the Inh cluster was consistently observed across the three ripple types. **E.** Activity centroid (i.e. center of mass of average activity in the ± 1 s surrounding a ripple) of each ensemble and thalamus around dCA1 or iCA1 ripples. **F.** Comparison of ripple-associated suppression of activity in thalamus and in the Inh ensemble between dCA1 and iCA1 ripples. Ripple-associated suppression was measured as the area under the curve during the period from -0.5 s to 0.5 s from ripple onset, analyzed for dCA1 and iCA1 ripples. We did not compare dAct and iAct ensembles here, because their activity must be biased toward either dCA1 or iCA1 ripples by definition.

**Fig. S2.**
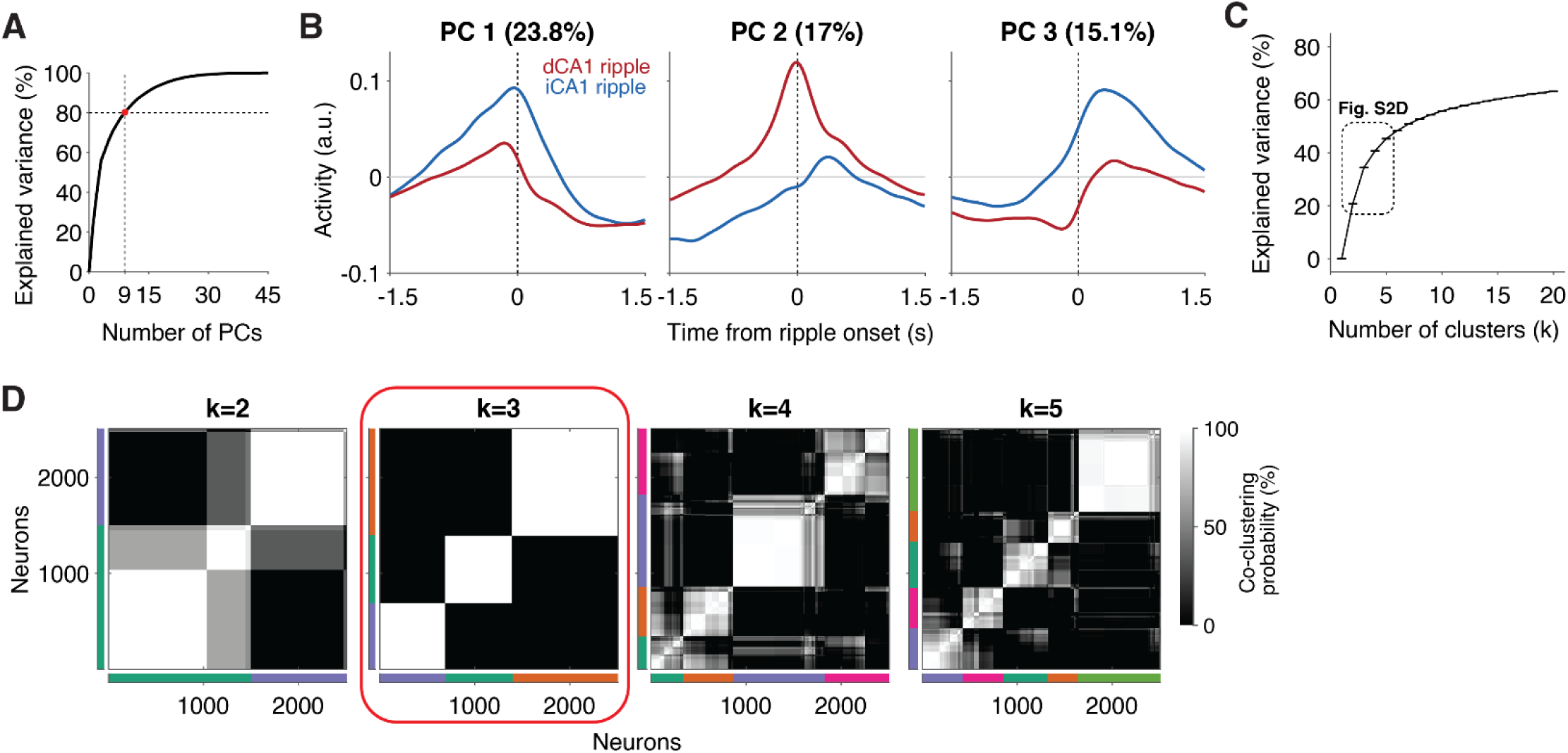
Details of k-means clustering. **A.** Explained variance of average neural activity profiles surrounding ripples as a function of the number of principal components (PCs). Principal component analysis (PCA) was performed using concatenated neural activity profiles surrounding dCA1 and iCA1 ripples (see Methods). The first 9 PCs, whose explained variance together reached 80.2 % (red dot), were used for k-means clustering. **B.** Activity profiles of the first three PCs around dCA1 and iCA1 ripples. Titles indicate % variance explained. **C.** Cumulative explained variance of the dataset composed of the first 9 PCs from A-B, as a function of the number of clusters used for k-means clustering. Please note that the graph has an elbow around k=2-5. **D.** Co-clustering probability of neurons from 100 iterations of k-means clustering for each k. This shows that the clustering result is most stable when k=3. Thus, for subsequent analyses, we classified ripple-coupled cortical neurons as belonging to one of three clusters.

**Fig. S3.**
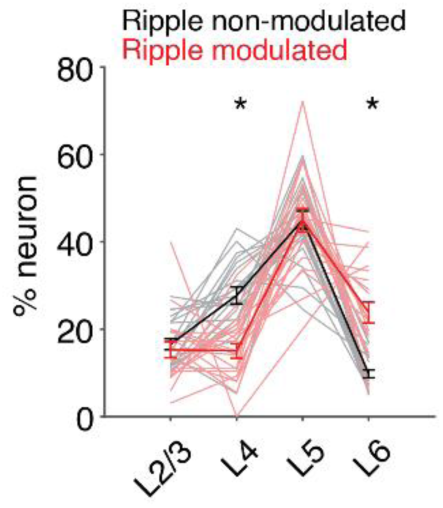
Layer distribution of each cluster. Fraction of ripple-modulated neurons and of non-modulated neurons across different layers of cortex. Ripple-modulated neurons were more located in layer 6, and less located in layer 4, in compared to non-modulated neurons. Each light gray or red line represents an individual session. Black and dark red lines: mean ± s.e.m.

**Fig. S4.**
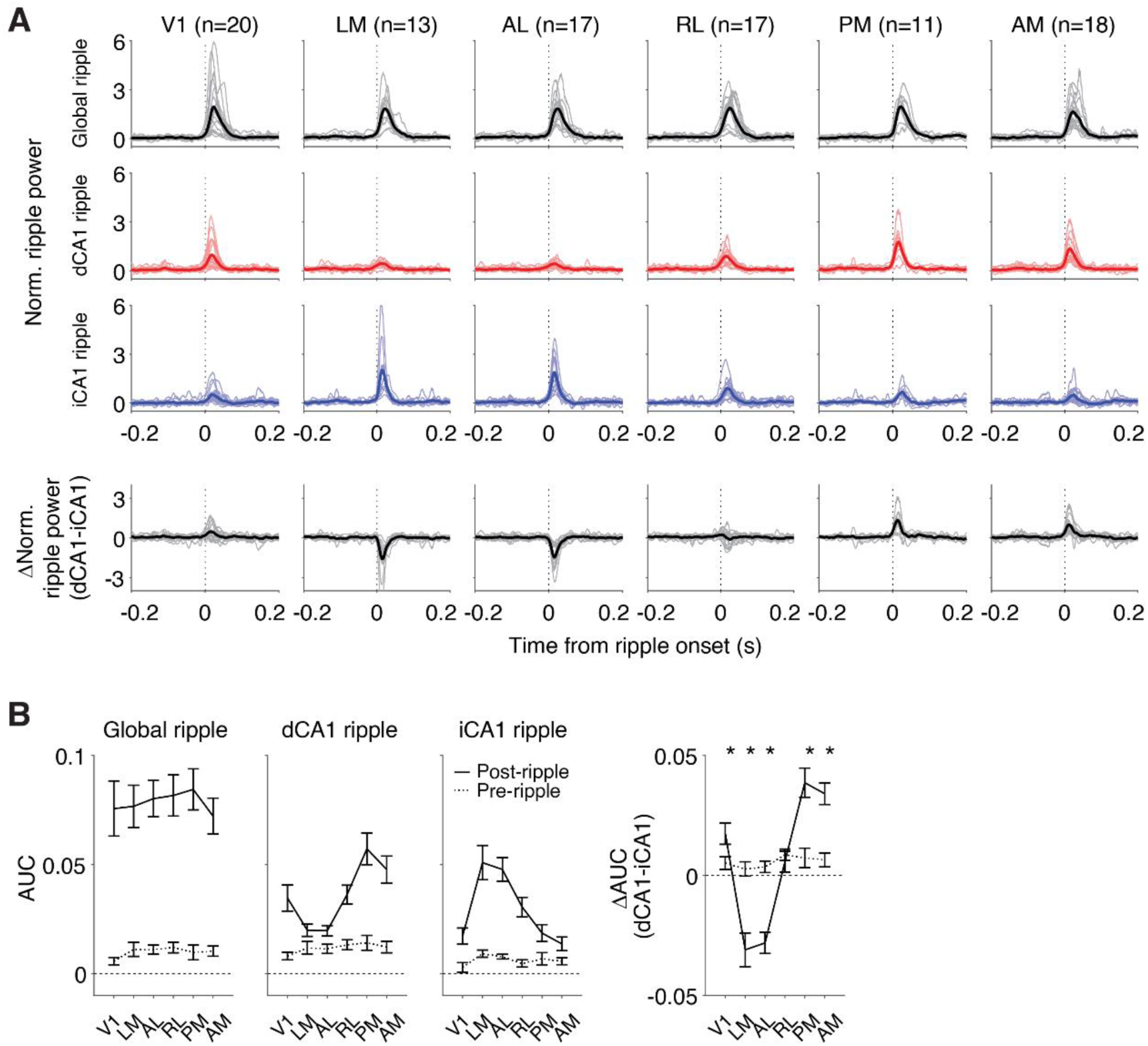
Ripple band power in medial and lateral visual cortex is differentially coupled to dCA1 and iCA1 ripples. **A.** Normalized power in the ripple band (100-250 Hz) in visual cortical areas in the period surrounding three different types of hippocampal ripples: global, dCA1 and iCA1 ripples. Around global ripples, ripple-band power similarly increased in all six visual cortical areas. By contrast, medial visual cortical areas (PM and AM) and lateral visual cortical areas (LM and AL) preferentially increased in ripple-band power around dCA1 and iCA1 ripples, respectively. Bottom row: difference in normalized cortical ripple-band power in each area during dCA1 vs. iCA1 ripples. V1, primary visual cortex; LM, lateromedial; AL, anterolateral; RL, rostrolateral; PM, posteromedial; AM, anteromedial. **B.** Left: area under the curve (AUC) during the 0.1 s window before (Pre-ripple) and after (Post-ripple) ripple onset. Right: difference in AUC for cortical ripple-band power during dCA1 vs. iCA1 ripples.

**Fig. S5.**
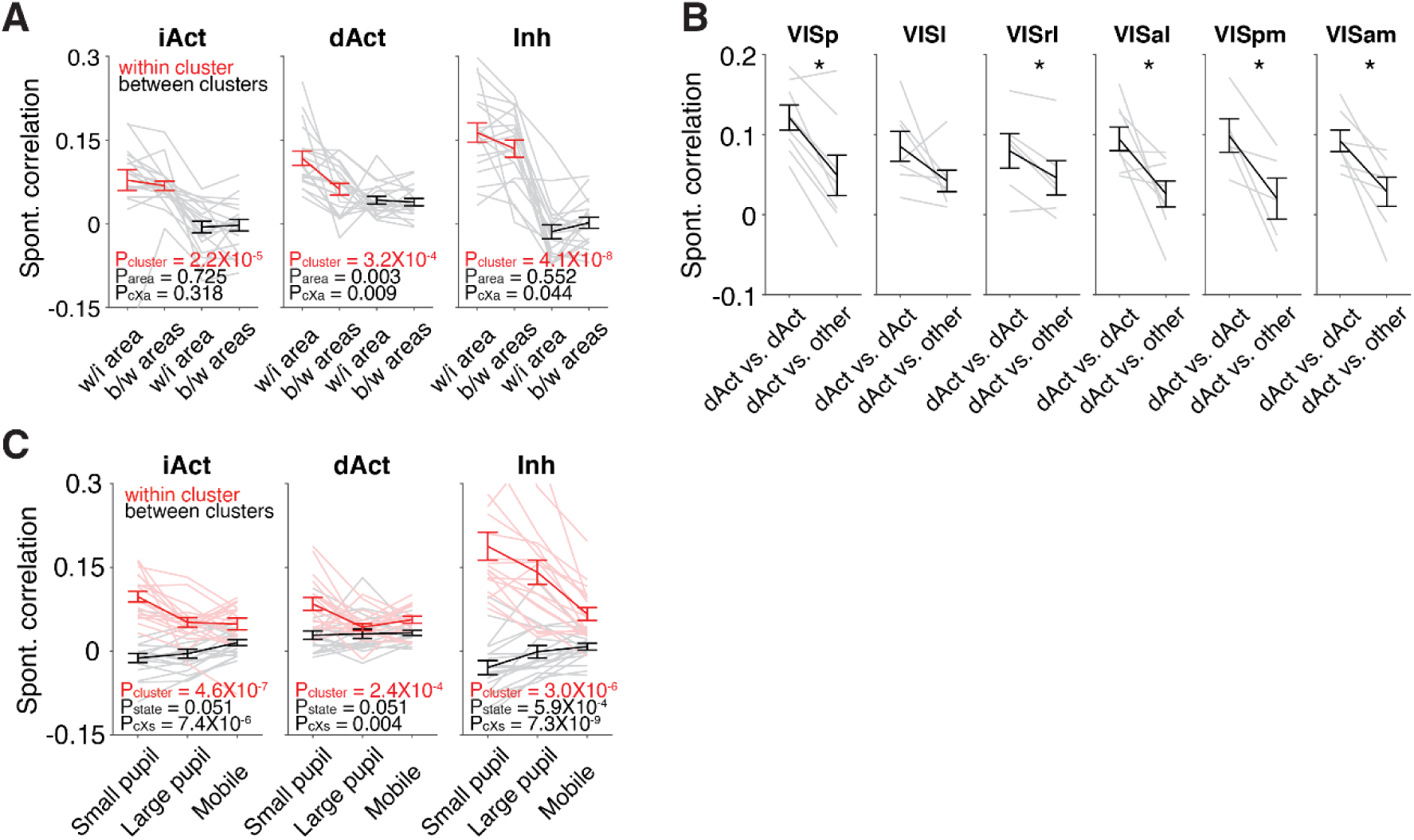
Each cluster represents a functional ensemble. **A.** Mean spontaneous neuronal correlations within (w/i area) and between areas (b/w areas) for each cluster. P_cluster_, P_area_, P_cxa_ are p-values for effect of cluster, area, and cluster × area interaction, respectively (two-way mixed ANOVA). Same as Fig. 2C, but each cluster was separately analyzed. **B.** Mean spontaneous correlations of dAct-dAct pairs (within cluster) and dAct-other cluster pairs (between clusters) in each visual area. For each area, only sessions in which there were more than 3 pairs each of dAct-dAct and dAct-other were used for analysis. **C.** Mean spontaneous correlations in three different behavioral states (small pupil, large pupil, and mobile). Same as Fig. 2H, but each cluster was separately analyzed. P_cluster_, P_state_, P_cxs_ are p-values for effect of cluster, state, and cluster × state interaction, respectively. For all clusters, even after accounting for whether the pairs were from the same or different areas (**A**) and behavioral state (**C**), spontaneous neural correlations for pairs within the same cluster were higher than correlations of pairs of neurons belonging to different clusters (P_cluster_, main effect of cluster).

**Fig. S6.**
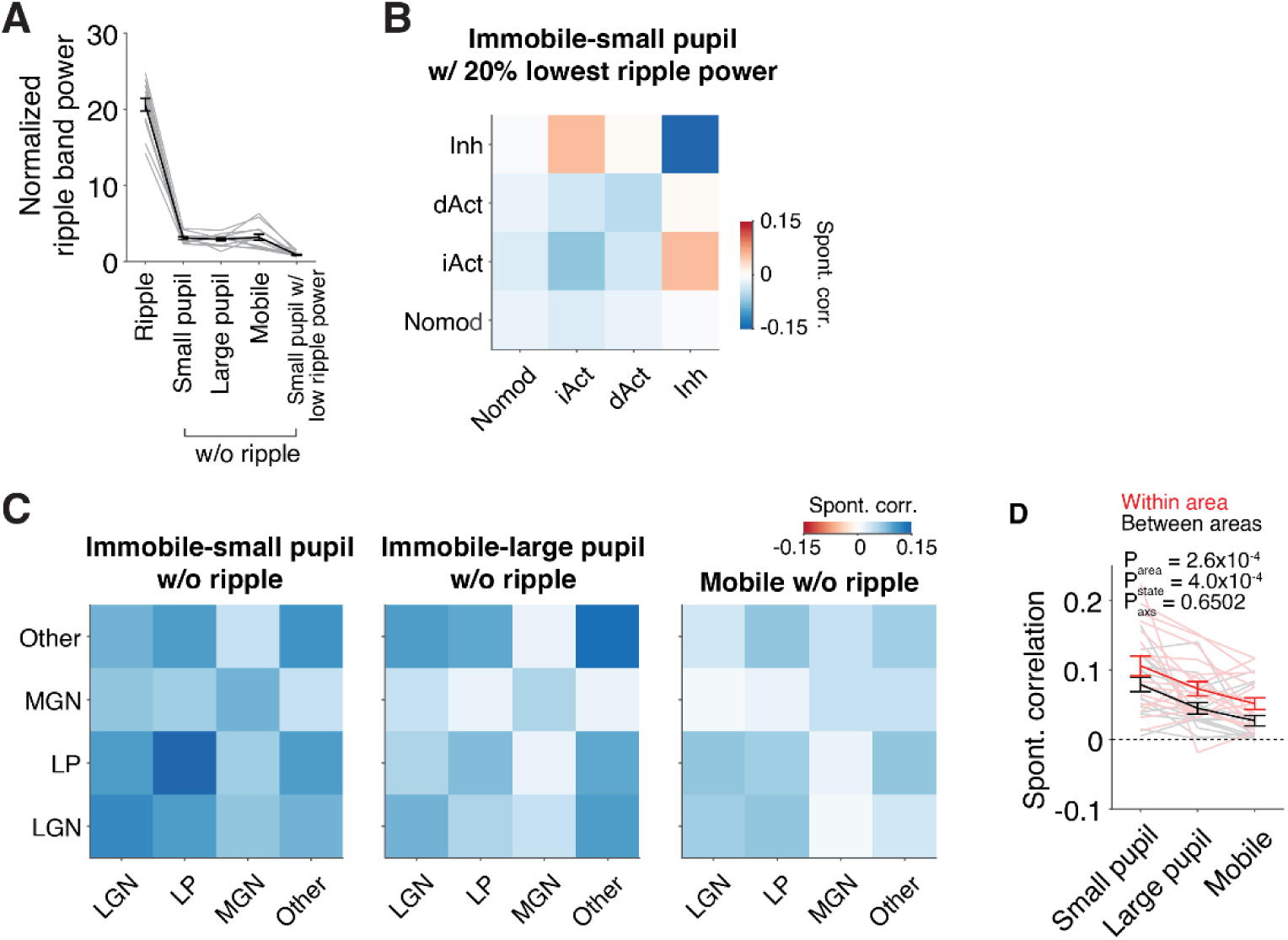
Additional analyses of spontaneous correlations. **A.** Normalized ripple band power (100-250 Hz) during different states. Ripple, small pupil, large pupil, and mobile states are the same states that are used in Fig. 2. Additionally, the subset of the non-ripple state with the 20% lowest ripple power and with small pupil size (below 50% of pupil size in the mobile state) was selected for analysis in **B**. **B.** Spontaneous correlation among clusters in visual cortex during small pupil state with low ripple power. The main properties of correlation structure, such as stronger correlation within cluster than between clusters and a negative correlation between the iAct and Inh ensemble, remained even in these moments with negligible ripple band power. **C.** Spontaneous correlation between thalamic neurons across visual (LP and LGN) and non-visual (MGN and Other) thalamic areas. Similar to Fig. 2G, correlations were separately calculated for three states with different levels of arousal during the spontaneous period. **D.** Average spontaneous correlation among neurons within the same area (Within area) or between different areas (Between areas). Spontaneous correlation was higher for a pair of neurons within area than across areas, but importantly, both within-area and between-areas correlations were significantly positive (t-test with Bonferroni correction, Within area, p<0.001 for all three states, Between area, p<0.05 for all three states).

**Fig. S7.**
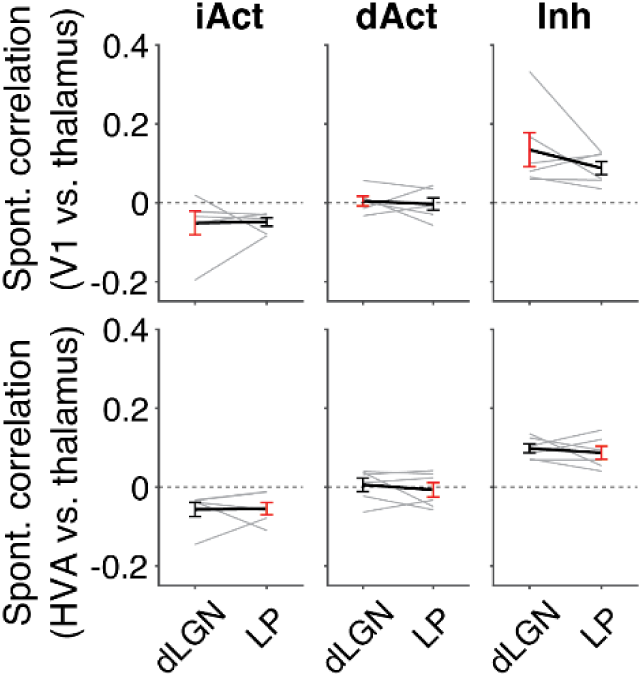
Spontaneous correlation of cortical ensembles with their main input thalamic area was not different from that with other thalamic area. Spontaneous correlation was measured between each ensemble from primary visual cortex (V1) and two visual thalamic areas, either dorsal lateral geniculate nucleus (dLGN) or lateral posterior nucleus (LP). The same analysis was done using ensembles in higher visual areas (HVA). V1 receives the main feedforward input from dLGN, while HVA receive stronger feedforward inputs from subregions of LP. Nevertheless, we could not find any significant difference in spontaneous correlation between dLGN-V1 pairs and LP-V1 pairs or between dLGN-HVA pairs and LP-HVA pairs.

## Methods

### Experimental data

We used an Allen Institute Neuropixels dataset that is freely available to the public (https://allensdk.readthedocs.io/en/latest/visual_coding_neuropixels.html). Specifically, we used the ‘Functional Connectivity’ dataset, which involves neural activity recorded from up to six Neuropixels probes while head-fixed mice were exposed to a variety of visual stimuli, including a 30-minute block of mean luminance gray screen. Analyses were conducted on this 30-minute block (‘spontaneous’ period), unless otherwise noted, in order to study coordinated neural dynamics across brain regions under conditions of minimal visual stimulation. The data were downloaded as NWB files and converted to MATLAB format before analysis. Animals with a total duration of immobility < 100 s during the spontaneous period (4/26 animals) or no LFP data (1/26 animal) were excluded from the analysis for clustering of visual cortical neurons according to ripple-associated responses. Additionally, we excluded one animal whose maximum distance between hippocampal recording sites was less than 2 mm in order to have sufficient distance separating the locations of ripple recordings. In total, 20 mice (16 males) were used for the clustering, including 13 wild-type (WT), two Pvalb-IRES-Cre/wt;Ai32(RCL-ChR2(H134R) EYFP)/wt, three Sst-IRES-Cre/wt;Ai32(RCL-ChR2(H134R) EYFP)/wt, and two Vip-IRES-Cre/wt;Ai32(RCL-ChR2(H134R) EYFP)/wt mice.

### Sharp-wave ripple detection and classification

LFPs were recorded from one fourth of all channels. We identified the CA1 pyramidal layer based on the density of single units recorded for each LFP channel near the hippocampus (the sum of all single units detected from five adjacent channels). LFPs from the channel with the most single units and the two adjacent channels were filtered using a 4^th^ order Butterworth band-pass filter (100-250Hz). Filtered LFP signals were squared, z-normalized, and averaged over the three channels to yield normalized ripple band power. We defined an event as a putative ripple if the ripple band power exceeded 2 SDs for more than 20 ms and the peak power exceeded 5 SDs during this time. If a subsequent ripple occurred less than 30 ms after the previous ripple ends, we merged them into a single ripple event. We excluded ripples that lasted longer than 250 ms. Ripples were included only during periods when the animal’s smoothed running speed (Gaussian filter, σ = 1 s) remained below 2 cm/s for more than 3 s.

Ripples in the dCA1 and iCA1 were detected using LFPs recorded from the most medially and most laterally located probes, respectively. We defined an event as a “global ripple” if two ripple events detected in dCA1 and iCA1 overlapped in time. The onset of the global ripple was determined as the onset of the earlier of the two ripple events. A ‘dCA1 ripple’ or a ‘iCA1 ripple’ was defined as a ripple that was detected only in dCA1 or iCA1, respectively (i.e., ripple power in the other region did not cross threshold for ripple detection).

### Visual cortex ripple band power

For each visual area, the normalized ripple band power was calculated from 5 channels that were randomly chosen. The averaged ripple band power was then aligned to hippocampal ripples. In this analysis, only those hippocampal ripples that had no other ripples within ±0.5 s of ripple onset were included. Ripple modulation of visual cortex ripple band power was quantified by calculating the area under the curve (AUC) of the ripple band power during the 0.1-s window before and after hippocampal ripple onset.

### Determining ripple-modulated neurons

A generalized linear model (GLM) was used to identify visual cortex neurons that were significantly modulated by dCA1 and/or iCA1 ripples during the spontaneous period:

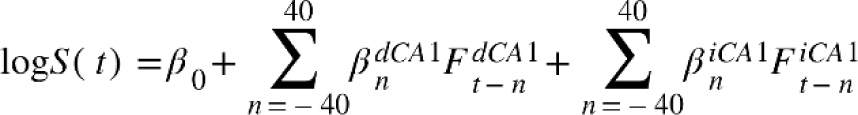

where S(t) is averaged firing rate during a 50-ms time bin, *β*_0_ to *β*_*n*_ are GLM coefficients, and *F_t−n_* is an occurrence of a ripple at time ( and represent dCA1 and iCA1 ripples, respectively; both global and local ripples were included in the analysis). A neuron was considered ‘ripple modulated’ if the GLM coefficients were significant in more than two consecutive bins within ±0.5 s of ripple onset. For the significance testing of % ripple modulated neurons between medial and lateral visual cortices (Fig. 1G, 1J), we excluded 3 out of 20 sessions lacking either medial or lateral visual cortex neurons.

### Clustering of visual cortical neurons

The averaged firing rates (50ms time bin, σ=50ms) during the [-1.5, 1.5] s time window since ripple onset were calculated separately for dCA1 and iCA1 ripples for all ripple-modulated neurons. The averaged firing rates around dCA1 and iCA1 ripples were concatenated and z-normalized for each neuron. This resulted in a matrix of size (number of neurons) × (number of time bins). Data dimensionality was reduced using PCA to (number of neurons) × (9), since the first 9 PCs explained >80% of the variance. We used *k* -means clustering with a wide range of values of *k* (2-20) to determine the optimal number of clusters (*k*). The percentage of variance explained was plotted as a function of *k*, and an ’elbow’ of the curve was sought (Fig. S2C). After estimating the range of *k* that sits around the elbow, we repeated *k* -means clustering 100 times for these putative *k*s, which ranged from 2 to 5. For each possible pair of neurons, the iterated clustering results were used to calculate the likelihood of being clustered in the same cluster (co-clustering probability; Fig. S2D). We found k = 3 clusters to be the optimal number, as this produced the highest likelihood of co-clustering.

### Spontaneous correlation

Each neuron’s spike counts during the spontaneous period were divided into 2-s time bins. Any time bin that contained a ripple event was excluded from this analysis. Pearson’s correlation was calculated for all possible neuronal pairs before being averaged across all neuronal pairs in the same cluster (Fig. 2B). One session with only five ripple-modulated neurons was excluded from this analysis. The mean number of neurons in each cluster (±sem across 19 sessions) was 26.37±3.29 for iAct, 41.74±10.74 for dAct, and 22.89±2.49 for Inh neurons.

Each time bin (2 s) was assigned to one of three states—mobile, immobile-large pupil, or immobile-small pupil— to estimate behavioral state-dependent spontaneous correlation. The mobile state was when the animal’s running speed exceeded 2 cm/s. The immobile-small pupil state was when the animal was immobile and the pupil size was smaller than 50% of the averaged pupil size during the mobile state, and the remaining immobile period was defined as the immobile-large pupil state. Spontaneous correlation at each behavioral state was calculated using spike count data associated with the corresponding behavioral state. To compare spontaneous correlation across behavioral states (Fig. 2G, H), we used only the sessions with the duration of each behavioral state >50 s (n = 18 sessions). The mean (±sem across 19 sessions) duration was 424 ± 53.13 s for immobile-small pupil state, 299.11 ± 41.58 s for immobile-large pupil state, and 537.11 ± 90.21 s for mobile state. In Fig. 4B, C, the analysis excluded one session in which no neurons were recorded from the thalamus (total 17 sessions). In Fig. S5B, the spontaneous correlation was calculated using only the time periods with the lowest 20% of ripple band power in the small pupil state (14 sessions with the duration of each behavioral state >50s). In Fig. S6, spontaneous correlation was separately calculated for V1-dLGN, V1-LP, HVA-dLGN, and HVA-LP pairs to see if the cortical ensemble has a higher correlation with the visual thalamic area providing the main feedforward input. Only sessions with both LP and dLGN neurons were used for this analysis (n = 10 sessions).

### State-dependent modulation

Running- and pupil-dependent modulation indices were calculated as the following:

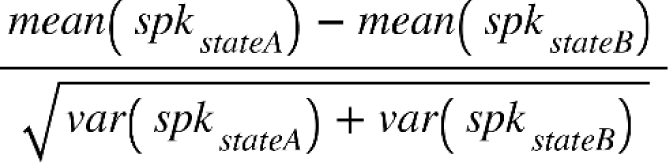

where *spk_stateA_* and *spk_stateB_* are 1-s binned spike counts during two different states, and . For the running-dependent modulation index, is the mobile state (speed > 2 cm/s) while is the immobile state. For the pupil-dependent modulation index, is the immobile state with top 25% of pupil sizes while is the immobile state with bottom 25% of pupil sizes.

### Analysis of visual properties

The size and significance of each neuron’s receptive field were determined based on neuronal responses to 81 locations of the Gabor stimulus, which were given at the beginning of each session in the Neuropixels data set (See Siegle et al., 2021 for details of receptive field analyses). Neurons with a significant receptive field had a receptive field size < 2500 degrees^2^ and a p-value<0.01 (bootstrapping with location-shuffled data). A neuron’s preferred stimulus condition was defined as the grating that elicited the greatest change in firing rate from the baseline period (0.5 s prior to grating onset) among eight different drifting gratings (4 orientations and 2 contrasts). We compared the firing rate during the 2-s grating presentation in each neuron’s preferred stimulus condition to that during baseline. A neuron was considered responsive to gratings if its firing rate was modulated significantly by grating presentation (p<0.05, paired t-test) and if it had a significant receptive field. Only grating-responsive neurons were analyzed in Fig. 3E-G. Each neuron’s responsiveness to its preferred grating stimulus was calculated as follows:

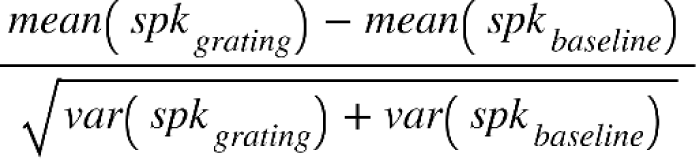

where *spk_grating_* and *spk_baseline_* are averaged firing rates during the 2 s after and the 0.5 s before the onset of the grating, respectively, in the preferred stimulus condition for each neuron.

Selectivity to the preferred grating stimulus was calculated as follows:

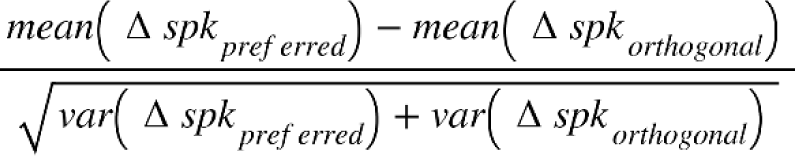

where Δ *spk_preferred_* and Δ *spk_orthogonal_* are baseline-subtracted firing rates during the preferred and orthogonal gratings, respectively, in the preferred contrast condition.

The phase modulation of a neuron reflects how the temporal frequency of a drifting grating stimulus modulates its activity. Response latency is defined as the time between the onset of the flash stimulus and the first spike. We used the estimates from the original study (Siegle et al., 2021) that were part of the Neuropixels data set. To compare the dependence of a neuron’s state-dependent modulation or visual tuning value on its cluster identity, we calculated the explained variance of each modulation index or visual tuning value by the cluster identity. The same analysis was run 100 times on neurons with randomly assigned cluster identities to calculate the explained variance under the null hypothesis.

### Cross-correlation analysis

Each neuron’s spike counts were divided into 20-ms bins, z-normalized, and averaged across neurons in the same cluster. Pearson’s correlation was calculated between the averaged firing rates of two clusters with time lags of up to ±1s. For a given session, cross-correlation was performed iteratively for all time windows longer than 3 s lacking a ripple event and then averaged over windows. Only sessions which of sum of all time windows in each behavior state is above 50s were used (n = 15 sessions). To determine whether the cross-correlogram is biased (i.e., whether the activity of one cluster leads the activity of the other cluster on average), the difference in the area under the curve (ΔAUC) was calculated during the 0.5 s window before and after a time lag of zero.

### Analysis of pre-ripple activity

To determine whether neural activity precedes a hippocampal ripple event, we calculated the time at which averaged ripple-associated neural activity reaches the centroid of activity during ±1 s from ripple onset. The averaged ripple-associated neural activity was normalized to the baseline (the activity between 3 s and 2 s before ripple onset). We then calculated the correlation between cortical neural activity 1-s prior to ripple onset and the strength of the subsequent ripple, as well as the correlation between hippocampal ripple band power 1-s prior to ripple onset and the strength of the subsequent ripple (as a control against possible predictability of ripple strength by pre-ripple cortical activity). The strength of a ripple was defined as the peak of normalized ripple band power (See <u>Sharp-wave ripple detection and classification)</u> within each ripple event. One session with only five ripple-modulated neurons and one session without thalamus recording were excluded from this analysis.

### Statistics

Details of statistical testing for each figure is shown in Table. S1.

**Table. S1.** Statistical results.

| Figure | Description | Test | Statistic | p value | Number of samples |
| --- | --- | --- | --- | --- | --- |
| Fig. 1G | % ripple modulated (medial vs. lateral visual cortices) | Paired t test | t = 3.22 | Two-tailed p value = 0.0053 | n = 17 sessions |
| Fig. 1I | Normalized depth | Kolmogorov-Smirnov test with Bonferroni correction | k =<br>Nomod vs. iAct, 0.26<br>Nomod vs. dAct, 0.21<br>Nomod vs. Inh, 0.28<br>iAct vs. dAct, 0.07<br>iAct vs. Inh, 0.09<br>dAct vs. Inh, 0.10 | Two-tailed p values =<br>Nomod vs. iAct, $6.0 \times 10^{-23}$<br>Nomod vs. dAct, $4.4 \times 10^{-21}$<br>Nomod vs. Inh, $3.6 \times 10^{-24}$<br>iAct vs. dAct, 0.7859<br>iAct vs. Inh, 0.2037<br>dAct vs. Inh, 0.0254 | n = 5358 neurons (all regular spiking neurons) |
| Fig. 1J | % cluster (medial vs. lateral visual cortices) | Paired t test | t =<br>iAct, 24.36<br>dAct, -19.49<br>Inh, -1.55 | Two-tailed p values =<br>iAct, $1.8 \times 10^{-61}$<br>dAct, $6.2 \times 10^{-48}$<br>Inh, 0.1227 | n = 17 sessions |
| Fig. 2B | Spontaneous correlation | Paired t test | t =<br>within vs. between, 9.42<br>within vs. nomod, 11.48 | Two-tailed p values =<br>Within vs. between, $2.3 \times 10^{-8}$<br>Within vs. nomod, $4.0 \times 10^{-10}$ | n = 19 sessions |
| Fig. 2C | Spontaneous correlation with an accounting for area difference | Repeated measures ANOVA | F =<br>Cluster, 86.82<br>Area, 1.81<br>Cluster x area, 5.20 | One-tailed p values =<br>Cluster, $2.6 \times 10^{-8}$<br>Area, 0.1955<br>Cluster x area, 0.035 | n = 19 sessions |
| Fig. 2E | Running modulation index | Two-way ANOVA | F =<br>Cluster, 69.19<br>Area, 3.05<br>Cluster x area, 2.58 | One-tailed p values =<br>Cluster, $8.7 \times 10^{-44}$<br>Area, 0.0094<br>Cluster x area, $7.3 \times 10^{-4}$ | n = 4769 neurons (regular spiking neurons in one of six visual cortices: V1, LM, AL, RL, PM, AM) |
| | Pupil modulation index | Two-way ANOVA | F =<br>Cluster, 177.16<br>Area, 5.38<br>Cluster x area, 2.88 | One-tailed p values =<br>Cluster, $7.1 \times 10^{-109}$<br>Area, $6.2 \times 10^{-5}$<br>Cluster x area, $1.6 \times 10^{-4}$ | |
| Fig 2F | Running speed across different states | Paired t-test with Bonferroni correction | t =<br>Ripple vs. low pupil, -3.18<br>Ripple vs. high pupil, -5.57<br>Ripple vs. Mobile, -5.55<br>Low pupil vs. high pupil, -4.68<br>Low pupil vs. Mobile, -5.54<br>High pupil vs. Mobile, -5.48 | Two-tailed p values =<br>Ripple vs. low pupil, 0.0329<br>Ripple vs. high pupil, $2.1 \times 10^{-4}$<br>Ripple vs. Mobile, $2.1 \times 10^{-4}$<br>Low pupil vs. high pupil, 0.0013<br>Low pupil vs. Mobile, $2.2 \times 10^{-4}$<br>High pupil vs. Mobile, $2.5 \times 10^{-4}$ | n = 4769 neurons (regular spiking neurons in one of six visual cortices: V1, LM, AL, RL, PM, AM) |
| | Normalized pupil size across different states | Paired t-test with Bonferroni correction | t =<br>Ripple vs. low pupil, 0.64<br>Ripple vs. high pupil, -12.65<br>Ripple vs. Mobile, -15.52<br>Low pupil vs. high pupil, -13.88<br>Low pupil vs. Mobile, -51.62<br>High pupil vs. Mobile, -13.46 | Two-tailed p values =<br>Ripple vs. low pupil, 1<br>Ripple vs. high pupil, $2.7 \times 10^{-9}$<br>Ripple vs. Mobile, $4.0 \times 10^{-11}$<br>Low pupil vs. high pupil, $6.4 \times 10^{-10}$<br>Low pupil vs. Mobile, $2.4 \times 10^{-19}$<br>High pupil vs. Mobile, $1.0 \times 10^{-9}$ | |
| | Normalized ripple band power across different states | Repeated measures of ANOVA | dCA1 probe<br>t =<br>Ripple vs. low pupil, 24.15<br>Ripple vs. high pupil, 23.56<br>Ripple vs. Mobile, 18.77<br>Low pupil vs. high pupil, 0.05<br>Low pupil vs. Mobile, -0.58<br>High pupil vs. Mobile, -0.82 | dCA1 probe<br>Two-tailed p values =<br>Ripple vs. low pupil, $8.1 \times 10^{-14}$<br>Ripple vs. high pupil, $1.2 \times 10^{-13}$<br>Ripple vs. Mobile, $5.0 \times 10^{-12}$<br>Low pupil vs. high pupil, 1<br>Low pupil vs. Mobile, 1<br>High pupil vs. Mobile, 1 | |
|  |  |  | iCA1 probe<br>t = | iCA1 probe<br>Two-tailed p values = |  |
| | | | Ripple vs. low pupil, 17.64<br>Ripple vs. high pupil, 18.95<br>Ripple vs. Mobile, 18.60<br>Low pupil vs. high pupil, 2.35<br>Low pupil vs. Mobile, 3.76<br>High pupil vs. Mobile, 1.49 | Ripple vs. low pupil, $1.4 \times 10^{-11}$<br>Ripple vs. high pupil, $4.3 \times 10^{-12}$<br>Ripple vs. Mobile, $5.9 \times 10^{-12}$<br>Low pupil vs. high pupil, 0.1873<br>Low pupil vs. Mobile, 0.0093<br>High pupil vs. Mobile, 0.9338 | |
| Fig 2H | Spontaneous correlation with an accounting for state difference | Repeated measures of ANOVA | F =<br>Cluster, 69.81<br>State, 8.03<br>Cluster x state, 58.34 | One-tailed p values =<br>Cluster, $2.0 \times 10^{-7}$<br>State, 0.001<br>Cluster x state, $1.0 \times 10^{-11}$ | n = 18 sessions |
| Fig 3A, left | Fraction of neurons having significant receptive field | Chi-square test | $\chi^2 = 21.12$ | One-tailed p value = $7.7 \times 10^{-4}$ | n = 5358 neurons (all regular spiking neurons) |
| | Fraction of neurons having significant receptive field without each group | Chi-square test following Bonferroni correction | $\chi^2$ =<br>w/o Nomod, 0.22<br>w/o iAct, 14.63<br>w/o dAct, 14.39<br>w/o Inh, 17.94 | One-tailed p values =<br>w/o Nomod, 1<br>w/o iAct, 0.022<br>w/o dAct, 0.025<br>w/o Inh, 0.005 | n =<br>w/o Nomod, 1734<br>w/o iAct, 4854<br>w/o dAct, 4565<br>w/o Inh, 4921 neurons |
| Fig 3A, right | Receptive field area | One-way ANOVA | F = 0.29 | One-tailed p values = 0.8311 | n = 3442 neurons (regular spiking neurons w/ significant receptive field) |
| Fig 3F | Responsiveness | Two-way ANOVA | F =<br>Cluster, 1.93<br>Area, 1.18<br>Cluster x area, 1.25 | One-tailed p values =<br>Cluster, 0.1226<br>Area, 0.3165<br>Cluster x area, 0.2296 | n = 3118 neurons (regular spiking neurons w/ significant receptive field in one of six visual areas: V1, LM, AL, RL, PM, AM) |
|  | Selectivity | Two-way ANOVA | F =<br>Cluster, 1.29<br>Area, 1.84<br>Cluster x area, 0.95 | One-tailed p values =<br>Cluster, 0.2774<br>Area, 0.1011<br>Cluster x area, 0.5031 |  |
| | Log10(phase modulation) | Two-way ANOVA | F =<br>Cluster, 0.15<br>Area, 33.15<br>Cluster x area, 0.83 | One-tailed p values =<br>Cluster, 0.9306<br>Area, $4.6 \times 10^{-33}$<br>Cluster x area, 0.6478 | |
|  | Response latency | Two-way ANOVA | F =<br>Cluster, 2.09<br>Area, 1.61<br>Cluster x area, 1.14 | One-tailed p values =<br>Cluster, 0.0989<br>Area, 0.1542<br>Cluster x area, 0.3108 |  |
| Fig 4E | Delta absolute AUC | Paired t test with Bonferroni correction | t =<br>Nomod, -0.49<br>iAct, 3.40<br>dAct, 1.94<br>Inh, -2.62 | Two-tailed p values =<br>Nomod, 1<br>iAct, 0.0171<br>dAct, 0.2936<br>Inh, 0.0804 | n = 15 sessions |
| Fig 4G | Delta absolute AUC | Paired t test with Bonferroni correction | t =<br>Nomod, 2.78<br>iAct, 4.17<br>dAct, 2.10 | Two-tailed p values =<br>Nomod, 0.0417,<br>iAct, 0.0025,<br>dAct, 0.1600 | n = 15 sessions |
|  | Difference of delta absolute AUC between Inh-Nomod and Inh-others | Paired t test with Bonferroni correction | T =<br>iAct, 3.31<br>dAct, 0.33 | Two-tailed p values =<br>iAct, 0.0095<br>dAct, 1 | n = 15 sessions |
| Fig 4I | Centroid time | t test | t =<br>Inh, -5.83<br>Thal, -2.60<br>iAct, -1.45<br>dAct, 2.91 | Two-tailed p values =<br>Inh, 2.0<br>Thal, 0.0187<br>iAct, 0.1658,<br>dAct, 0.0098 | n = 18 sessions |
| Fig 4J | Correlation between pre-ripple firing rate and ripple strength | t test | t =<br>Inh, -2.38<br>Thal, -3.18<br>iAct, 2.31<br>dAct, -0.54 | Two-tailed p values =<br>Inh, 0.029<br>Thal, 0.006<br>iAct, 0.034<br>dAct, 0.593 | n = 18 sessions |
| Fig 4K | Centroid time | t test | t =<br>Inh, -5.83<br>Thal, -2.6<br>iAct, -1.4<br>dAct, 2.9 | Two-tailed p values =<br>Inh, $2.0 \times 10^{-5}$<br>Thal, 0.0187<br>iAct, 0.1658<br>dAct, 0.0098 | n = 18 sessions |
| Fig. S1A, left | Ripple duration | Paired t test with Bonferroni correction | t =<br>dCA1 vs. iCA1, 5.35<br>dCA1 vs. global, -11.76<br>iCA1 vs. global, -15.94 | Two-tailed p values =<br>dCA1 vs. iCA1, $2.2 \times 10^{-4}$<br>dCA1 vs. global, $2.2 \times 10^{-9}$<br>iCA1 vs. global, $1.1 \times 10^{-11}$ | n = 20 sessions |
| Fig. S1A, right | Ripple strength | Paired t test with Bonferroni correction | t =<br>dCA1 vs. iCA1, -1.57<br>dCA1 vs. global, -7.78<br>iCA1 vs. global, -5.77 | Two-tailed p values =<br>dCA1 vs. iCA1, 0.8040<br>dCA1 vs. global, $1.5 \times 10^{-6}$<br>iCA1 vs. global, $8.8 \times 10^{-5}$ | n = 20 sessions |
| Fig. S1B | % global ripple | t test (vs. 50%) | t = 0.47 | Two-tailed p value = 0.6448 | n = 20 sessions |
| Fig. S1E | Centroid time | t test | t =<br>Inh, dCA1, -2.6<br>Thal, dCA1, -5.2<br>iAct, dCA1, -3.1<br>dAct, dCA1, 1.1<br>Inh, iCA1, -2.4<br>Thal, iCA1, -1.4<br>iAct, iCA1, -3.2<br>dAct, iCA1, 4.2 | Two-tailed p values =<br>Inh, dCA1, 0.0164<br>Thal, dCA1, $7.2 \times 10^{-5}$<br>iAct, dCA1, 0.0065<br>dAct, dCA1, 0.3077<br>Inh, iCA1, 0.0257<br>Thal, iCA1, 0.1845<br>iAct, iCA1, 0.0057<br>dAct, iCA1, $5.8 \times 10^{-4}$ | n = 18 sessions |
| Fig. S1F | Ripple-associated suppression | Paired t test | t =<br>Thalamus, 2.69<br>Inh, 5.24 | Two-tailed p values,<br>Thalamus, 0.0154<br>Inh, $6.7 \times 10^{-5}$ | n = 18 sessions |
| Fig. S3 | % neuron | Paired t test with Bonferroni correction | t =<br>L2/3, 0.56<br>L4, 5.59<br>L5, 0.02<br>L6, -5.68 | Two-tailed p values,<br>L2/3, 1<br>L4, $8.7 \times 10^{-5}$<br>L5, 1<br>L6, $7.2 \times 10^{-5}$ | n = 20 sessions |
| Fig. S4B, right | ΔAUC (dCA1-iCA1) | Paired t test with Bonferroni correction | t =<br>V1, -3.26<br>LM, 4.90<br>AL, 8.44<br>RL, 0.81<br>PM, -6.05<br>AM, -7.20 | Two-tailed p values =<br>V1, 0.0246<br>LM, 0.0022<br>AL, $1.6 \times 10^{-6}$<br>RL, 1<br>PM, $7.4 \times 10^{-4}$<br>AM, $9.0 \times 10^{-6}$ | n =<br>V1, 20 sessions<br>LM, 13 sessions<br>AL, 17 sessions<br>RL, 17 sessions<br>PM, 11 sessions<br>AM, 18 sessions |
| Fig. S5A | Spontaneous correlation | Two-way mixed ANOVA | F =<br>iAct:<br>Cluster, 32.18<br>Area, 0.13<br>Cluster x area, 1.06<br>dAct :<br>Cluster, 19.69<br>Area, 12.30<br>Cluster x area, 8.55<br>Inh:<br>Cluster, 81.87<br>Area, 0.37<br>Cluster x area, 4.70 | One-tailed p values =<br>iAct:<br>Cluster, $2.2 \times 10^{-5}$<br>Area, 0.7246<br>Cluster x area, 0.3179<br>dAct :<br>Cluster, $3.2 \times 10^{-4}$<br>Area, 0.0025<br>Cluster x area, 0.0091<br>Inh:<br>Cluster, $4.1 \times 10^{-8}$<br>Area, 0.5517<br>Cluster x area, 0.0438 | n = 19 sessions |
| Fig. S5B | Spontaneous correlation | Paired t test | t =<br>V1, 4.98<br>LM, 1.76<br>RL, 3.30<br>AL, 2.86<br>PM, 3.15<br>AM, 3.31 | Two-tailed p values =<br>V1, 0.0016<br>LM, 0.1293<br>RL, 0.0214<br>AL, 0.0243<br>PM, 0.0345<br>AM, 0.0162 | n =<br>V1, 8 sessions<br>LM, 7 sessions<br>RL, 6 sessions<br>AL, 8 sessions<br>PM, 5 sessions<br>AM, 7 sessions |
| Fig. S5C | Spontaneous correlation | Two-way mixed ANOVA | F =<br>iAct:<br>Cluster, 61.80<br>State, 3.26<br>Cluster x state, 17.07<br>dAct :<br>Cluster, 21.51<br>State, 3.25<br>Cluster x state, 6.65 | One-tailed p values =<br>iAct:<br>Cluster, $4.6 \times 10^{-7}$<br>State, 0.0508<br>Cluster x state, $7.4 \times 10^{-6}$<br>dAct :<br>Cluster, $2.4 \times 10^{-4}$<br>State, 0.0510<br>Cluster x state, 0.0037 | n = 18 sessions |
| | | | Inh:<br>Cluster, 46.36<br>State, 9.33<br>Cluster x state, 34.17 | Inh:<br>Cluster, $3.0 \times 10^{-6}$<br>State, $5.9 \times 10^{-4}$<br>Cluster x state, $7.3 \times 10^{-9}$ | |
| Fig. S6D | Spontaneous correlation | t test following Bonferroni correction | t =<br>Within area<br>Low pupil, 6.86<br>High pupil, 5.92<br>Mobile, 5.36<br>Between areas,<br>Low pupil, 6.64<br>High pupil, 4.79<br>Mobile, 3.13 | Two-tailed p values =<br>Within area<br>Low pupil, $3.1 \times 10^{-5}$<br>High pupil, $1.1 \times 10^{-4}$<br>Mobile, $3.0 \times 10^{-4}$<br>Between areas,<br>Low pupil, $3.3 \times 10^{-5}$<br>High pupil, $8.6 \times 10^{-4}$<br>Mobile, 0.0222 | n = 15 sessions |
| Fig. S7 | Spontaneous correlation | Paired t test | t =<br>V1, iAct, -0.09<br>V1, dAct, -0.18<br>V1, Inh, 1.72<br>HVA, iAct, -0.64<br>HVA, dAct, 0.58<br>HVA, Inh, 1.21 | Two-tailed p values =<br>V1, iAct, 0.9316<br>V1, dAct, 0.8607<br>V1, Inh, 0.1366<br>HVA, iAct, 0.5435<br>HVA, dAct, 0.5846<br>HVA, Inh, 0.2703 | n = 7 sessions |

